# Mast Cells Enhance Myeloma Engraftment and Promote Bone Destruction in the NSG-hIL6 Patient Derived Xenograft Model

**DOI:** 10.64898/2026.05.14.725220

**Authors:** Zainul S. Hasanali, Alfred L. Garfall, Dan T. Vogl, Adam D. Cohen, Adam J. Waxman, Sandra P. Susanibar-Adaniya, Shivani Kapur, Edward A. Stadtmauer, Cara Cipriano, Kristy Weber, David Allman

## Abstract

Multiple myeloma remains a fatal, incurable disease. Most therapies are targeted to the cancer cell or T cell engagement. Little is known about the supporting myeloma microenvironment and its contribution to tumor fitness. Here, we expand upon the observation of human mast cells in the NSG-hIL6 myeloma patient derived xenograft mouse model to show mast cells decrease time to engraftment, promote increased myeloma engraftment and cause myeloma bone disease. We identify 10 mast cell secreted factors that together improve the survival of patient myeloma cells *in vitro*. Our results highlight the versatility of the NSG-hIL6 model to study microenvironmental interactions between human bone marrow cells and myeloma and confirm prior suggestions that clinical signs of disease, such as osteolytic lesions, may at least partially be related to non-malignant bone marrow microenvironmental cells, such as mast cells.

## Introduction

Multiple myeloma (MM) is responsible for ∼100,000 deaths/year worldwide.(1) There are several efficacious therapies, including bispecific antibodies and chimeric antigen receptor T cells, but none are curative. Durability remains a significant issue in MM and cancer at large. In MM, malignant cells in the bone marrow, like their normal plasma cell counterparts, live in protected spaces, niches, that may allow them to survive MM targeted therapies.(2) Studies of the tumor microenvironment have been gaining momentum, but the complex interplays of non-cancer cells in tumors, particularly in MM, have only begun to be explored.

A major reason that studying the MM microenvironment has been difficult is the lack of appropriate models to dissect interplay between malignant and non-malignant cells. We recently published a novel patient derived xenograft (PDX) model using the NSG-hIL6 mouse(3), an immunodeficient mouse that expresses human IL6 and allows for the engraftment of all plasma cell dyscrasias, including MM. Over time we also noted the presence of MM clinical sequelae in PDX mice, including bone lesions. We further assessed the engrafted bone marrow in the PDX with scRNA sequencing and noted the presence of three populations of human cells, myeloma cells, exhausted T cells and surprisingly, mast cells.(3) The current study expands upon the presence of mast cells and seeks to explore their role in supporting MM engraftment and its effect on bone destruction.

Prior studies looking at histological mast cell density in core biopsies from MM patients revealed a correlation between mast cell density and ISS stage and therefore prognosis of disease.(4) Furthermore, analysis of patient bone marrow mast cell density correlated with osteolysis and is suggestive of a potential role of mast cells in myeloma bone disease.(5) In this study, we explore the ability of microenvironmental normal, non-malignant mast cells to support the engraftment of mouse and human myeloma in transplant and PDX settings, link their presence to myeloma bone disease and identify and validate a small number of mast cell factors that may be responsible for these observations.

## Materials and Methods

### scRNA sequencing

Single cell RNA sequencing data was downloaded from GEO accession numbers GSE246140, GSM7857100 and GSM785710 and analyzed with the Parse Biosciences Pipeline in Trailmaker.(3)

### Immunohistochemistry

All IHC stains were performed from 10% formalin fixed, EDTA decalcified, paraffin embedded 1μm sections at histowiz: histopathology. Antibodies used were CD138 (hu monkey) (Catalog # GA64261-2) and c-kit (Catalog # ab32363).

### Vκ*-myc transplant model

Vκ*-myc cells were obtained from the laboratory of Leif Bergsagel (Arizona). Frozen cells were thawed and resuspended in PBS prior to injection into either Jaxboy (C57BL/6J-Ptprc^em6Lutzy^/J – Jax 033076, RRID:IMSR_JAX:033076) or Sash mice (B6.Cg-Kit^W-sh^/HNihrJaeBsmJ – Jax 030764, RRID:IMSR_JAX:030764) retro-orbitally. Sera were collected weekly for 4 weeks and assessed for M spike. SPEP gels were scanned using a Biorad GelDoc EZ and band densities calculated with ImageLab (BioRad). Mice were euthanized at 4 weeks and flow cytometry performed for the presence of Vκ*-myc cells by intracellular kappa light chain staining.

### Mast cell generation

Basic Protocol #1 from “Generation, Isolation and Maintenance of Human Mast Cells and Mast Cell Lines”(6) was used to generate all mast cells. Deidentified CD34+ stem cells were obtained from the UPenn Stem Cell Laboratory as autologous stem cell transplant leuko packs from deceased patients. Mast cells were cultured for 10 weeks prior to use and assessed for the presence of FcERI (BioLegend 134324), CD117 (Kit) (BioLegend 332204) dual positivity prior to use (Supplemental figure 3).

### NSG-hIL6 patient derived xenograft model and analysis

Mice can be purchased from Jackson Laboratory (Strain #:028655). Detailed methods for myeloma sample preparation, set up of myeloma in the NSG-hIL6 model, CT scanning, assay of sera by ELISA for serum immunoglobulins and serum protein electrophoresis can be found in our previous study.(3) Deviations from our prior published protocol are as follows. Fresh cells were subjected to CD138+ bead selection (Miltenyi 130-111-744) on LS columns (Miltenyi 30-042-401) according to manufacturer instructions and checked for purity by flow cytometry using singlet, live/dead (Thermo L10119) and intracellular light chain staining (Supplemental Figure 2). Cells were injected into the left femur at a constant cell count, CD138+ alone (2×10^6^ per mouse), CD138+ with mast cells (each at 1×10^6^ cells per mouse) or mast cells alone (2×10^6^ cells per mouse).

### Cytokine profiling

Supernatants from MM90-mast cell co-culture from Figure 4A were applied to the R&D Systems Proteome Profiler Human XL cytokine array kit (ARY022B) per manufacturer’s instructions. Chemiluminescence was recorded on an Amersham Imager 600 and photon density analyzed with Image Lab (BioRad). Heatmaps were produced using GraphPad Prism. Complete table of cytokines tested is available in supplemental table 1.

### Cell culture

Myeloma mononuclear bone marrow patient samples were thawed from 10% DMSO in FBS frozen vials and placed into RPMI 1640 medium with 10% FBS and Glutamax (Thermo 35050061) in 1mL in 48 well tissue culture plates at a total cell count of 2×10^6^ cells per well. Cell counts were kept constant, so conditions of mixed myeloma and mast cells contained 1×10^6^ of each. For cytokine culture experiments, cytokines were added at the same time as myeloma cell plating at a dose of 200ng/mL. Cytokines were obtained from PeproTech (sCD14 110-01-10ug, Osteopontin 120-35-10ug, Dkk-1 120-30-2ug, CXCL5 300-22-5ug, GDF-15 120-28C-5ug, IGF-BP2 350-06B-5ug, IL6 200-06-20ug, Mip-3b 300-29B-20ug) or BioLegend (uPAR 559702, DPPIV 764102). A cell culture insert in a 24 well plate (Millipore PTHT06H48) was used for (Figure 4B). All cells were cultured for 5 days and then assessed for the presence of myeloma cells by flow cytometry as described.

### Flow cytometry

Cells were isolated from the left, injected femur from PDX mice and the left femur of Vκ*-myc transplanted mice. Cells were lysed for red blood cells using ACK lysis buffer for 5 minutes at room temperature and stained for live cells with zombie near IR live/dead (Thermo L10119) (10 minutes) and fluorescently labeled antibodies of markers of interest (30 minutes) in 0.1%BSA PBS buffer. To identify myeloma cells, cells were gated on singlets, live cells, CD3- (BioLegend 300323), CD20- (BioLegend 302311) and intracellular positive staining for lambda (Southern Biotech 2060-31) or kappa light chain (Southern Biotech 2060-02). Intracellular staining was carried out per manufacturer’s instructions using 10% formalin fixation for 10 minutes and PhosFlow Perm Buffer III (BD 558050).

### Study Approval

Informed consent was obtained prior to use of all patient samples per approved IRB protocol # 842940 at the University of Pennsylvania. No identifying information was accessed or used. Inclusion/exclusion criteria for myeloma samples used were based on frozen cell availability. Samples with at least 1e6 cells were used. There was no attrition. Sex of donors/age was not relevant to samples selection. No randomization/blinding/power analyses were relevant to this study. The stem cell and xenograft core IACUC protocol for animal model development was used for NSG-hIL6 experiments.

### Statistics and graphing

2-sided ANOVA and t-tests were used for comparison of >2 groups or 2 groups respectively and derived from GraphPad Prism. Significance cut offs were α=0.05. All error bars represent mean and standard deviation. p-values are listed on graphs. No statistics were attempted for groups with less than 3 replicates. All graphs were generated with GraphPad Prism (RRID:SCR_002798).

### Data Availability

scRNAseq data can be obtained from GEO as noted in the scRNAseq materials and method section. All other data generated in this study are available upon request from the corresponding author.

## Results

### Human mast cells are present in human myeloma engrafted NSG-hIL6 PDX mice

NSG-hIL6 mice were injected with human bone marrow mononuclear cells, including myeloma cells. Upon scRNA sequencing of 20,000 cells from human engrafted mouse marrow we noted three distinct populations of human cells by RNA expression profile, myeloma cells, T cells and mast cells. The finding of mast cells was unexpected.(3) To ensure our cell population of interest was in fact mast cells and not basophils or eosinophils which share overlapping transcriptional profiles, we analyzed a set of 11 genes in our scRNA dataset whose expression profile is highly specific for mast cells over other inflammatory cells (Figure 1A). This 11 gene signature was determined through prior analysis of allergic asthma inflammatory scRNA datasets.(7) Our scRNA seq dataset came from the marrow of a mouse engrafted with new diagnosis, untreated myeloma containing the t(4;14) translocation. To determine if this finding was unique to this myeloma patient sample or more generalizable, we performed immunohistochemistry staining on bone marrow from PDX mice engrafted with an additional 2 new diagnosis or 3 relapsed/refractory myeloma samples and noted human mast cells in all marrow samples (Supplemental Figure 1). Representative histology is shown in Figure 1B. Mast cells were identified with kit staining and toluidine blue staining (purple cells). Though kit expression is present in hematopoietic stem cells and some myeloma cells as well(8), the combination of toluidine blue positivity and the expression profile from scRNAseq supports mast cell identity. These data suggest an overrepresentation of mast cells in myeloma xenografts in hIL6 expressing mice.

**Fig 1:**
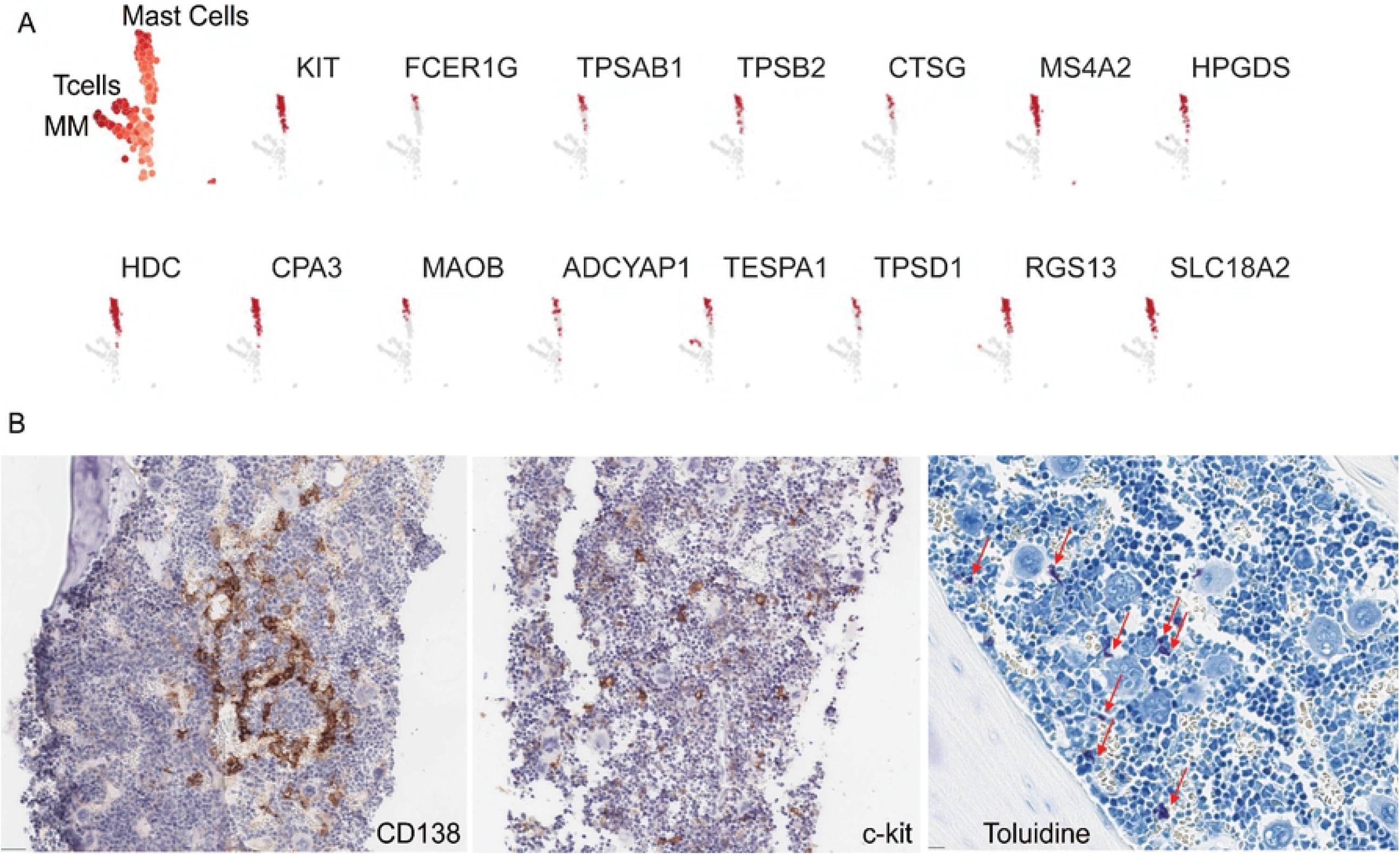
Human mast cells were overrepresented in scRNA sequencing in the NSG-hIL6 MM PDX model.

We reanalyzed single cell RNA sequencing of bone marrow from an NSG-hIL6 mouse intraosseously xenografted with mononuclear cells from the bone marrow of a new diagnosis multiple myeloma patient that was published previously(3). Within the subset of human cells, we previously identified 3 populations of human cells, one of which appeared to be a population of mast cells. We further analyzed our data set using an 11 gene expression signature that strongly predicted mast cell identity.(7) (A) We noted that all 11 genes, in addition to two canonical mast cell genes, KIT(CD117) and FCER1G(FcE receptor subunit G), were expressed nearly exclusively in the presumed mast cell group. (B) Immunohistochemistry staining of one PDX graft (MM93) representative of 2 new diagnosis myeloma patients and 3 multiply relapsed refractory patients showed the presence of numerous human kit+ cells in the bone marrow. CD138+ myeloma cells (left) and c-kit+ mast cells (middle). Toluidine blue (right) denotes mast cells (red arrows) by purplish appearance.

### Mast cell deficient C57BL/6 mice were less permissive and slower to engraft Vκ*-myc cells

After confirmation of human mast cells in the PDX model, we asked whether their presence represented a biological advantage for myeloma cells or were simply an artifact of the hIL-6 expressing NSG-hIL6 mouse given that IL6 is a required mast cell survival factor.(9) We obtained 8-week old Jaxboy (C57BL/6 mice congenic for the CD45.1 allele) or Sash (Kit mutated such that hematopoiesis is unaffected other than the absence of mast cells)(10-12) mice. Jaxboy mice were used as controls because of their availability in the mouse colony at the correct age. Vκ*-myc cells were retro-orbitally injected at 10^6^ per mouse (n=3 per group). Vκ*-myc cells are a mouse myeloma derived from the Vκ*-myc mouse model that are transplantable in C57BL/6 mice.(13) We collected sera weekly for 4 weeks and assessed for M-spikes representative of the Vκ*-myc graft growth by serum protein electrophoresis (SPEP). M-spikes rose significantly faster in Jaxboy mice vs sash mice (Figure 2A,B). The mice rapidly died after 4 weeks. Upon repeat with increased numbers (n=4), bone marrow flow cytometry showed increased myeloma cells in jaxboy vs sash mice (Figure 2C,D). These results support a biological role of mast cells in supporting myeloma cells *in vivo*.

**Fig 2:**
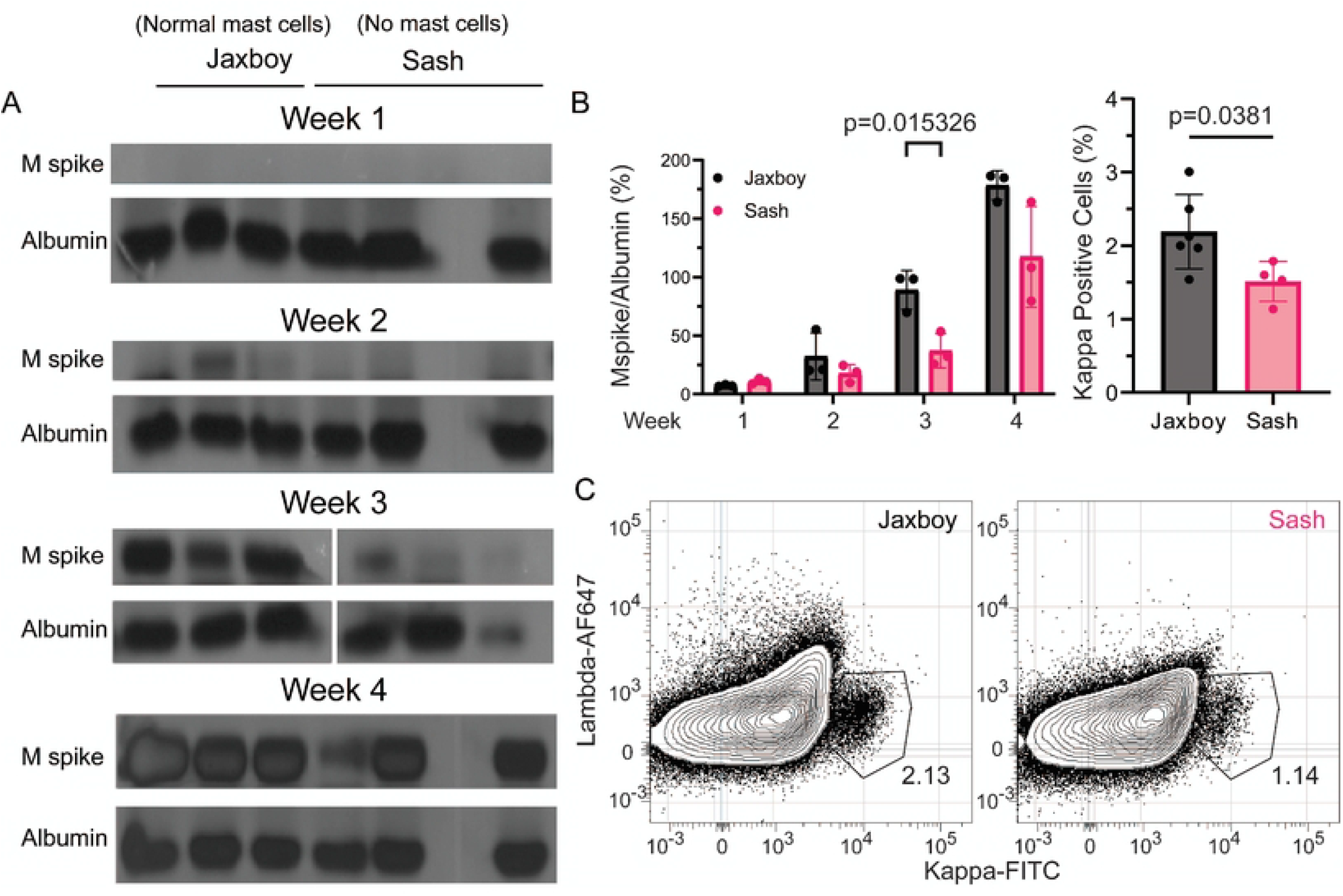
Mast cell deficient C57BL/6 mice engraft Vκ*-myc cells Slower than mast cell sufficient mice.

Three 8 week old Jaxboy (CD45.1 congenic B6) or Sash (mast cell deficient B6) mice were injected with 10^6^ Vκ*-myc cells retroorbitally to set up the Vκ*-myc transplantation model. (A) Blood was collected weekly for 4 weeks, serum separated with centrifugation and run on serum protein electrophoresis gels to quantitate M-spikes. Albumin bands were used as loading controls. (B) M-spike/albumin ratios were calculated from band intensities derived from the ImageLab software suite and plotted versus time. (C) Flow cytometry of kappa positive cells representative of Vκ*-myc cells in bone marrow of Jaxboy or Sash (no mast cells) mice. Cells are gated on singlet, lymphocyte region, live cells and intracellular kappa vs lambda staining. Technically because Vκ*-myc cells are CD45.2 and the Jaxboy mice are CD45.2, we could have gated on only CD45.2 positive cells. However, Sash mice were on a CD45.2 (usual C57BL/6 CD45 allele), so in order to compare between the two groups, total kappa positive cells were used rather than subtyped by CD45 allele in the Jaxboy vs total in the Sash. (D) Aggregate plot of kappa positive cells from bone marrow of Vκ*-myc Jaxboy (n=6) vs Sash (n=4) transplanted mice. p-values determined by student’s t-test.

### Co-injection of human mast and myeloma cells in NSG-hIL6 mice led to faster engraftment, increased myeloma burden and increased bone destruction

Though our Vκ*-myc transplant experiments were suggestive of a role for mast cells in supporting myeloma engraftment, they suffered from two problems. Firstly, the entire system was murine, casting some doubt on the relevance to human disease, and secondly, though well published, we were unable to detect the presence of mast cells in the bone marrow by flow cytometry in either the jaxboy or sash mice. This latter observation was likely due to the rarity of mast cells overall in normal biology and methodological limitations. Nonetheless, we sought an additional approach. We engrafted NSG-hIL6 mice (n=4 per group) with either human mast cells, 88% pure CD138+ bead selected patient myeloma cells (supplemental figure 2) from a new diagnosis kappa light chain myeloma patient who presented with impending tibial fracture (MM80), or both. Mast cells were generated *in vitro* from CD34+ stem cells from deidentified stem cell collections from deceased non-myeloma patients and phenotyped for the presence of FcE receptor and kit expression by flow cytometry prior to use using an established protocol (supplemental figure 3).(6) Sera were collected at regular intervals and ELISA performed for total immunoglobulin. These mice have no immunoglobulins at baseline, so all levels must come from the graft. In our characterization of the NSG-hIL6 model, we showed that detectable paraprotein matched that of the patient myeloma clone(3), so for simplicity, we chose to capture total immunoglobulins for these studies. Mast cell engrafted mice showed no antibody levels as expected. MM80 myeloma cells showed engraftment by week 28, but co-injected mice showed engraftment by week 15 (Figure 3A). These results were in line with those observed in Vκ*-myc transplantation experiments (Figure 2). Only 2 mice from each of the MM80 groups survived for bone marrow and CT scan analysis, thus statistics are not calculated for marrow flow cytometry. Mice were taken down at 35 weeks post cell injection. Upon bone marrow examination of the presence of light chain positive myeloma cells, mast cell engrafted mice had no appreciable light chain signal. MM80 had ∼1%, but MM80+mast cells had ∼3% (Figure 3B). There were still a small number of human mast cells detectable in the bone marrow in the mast cell alone condition and none in the MM80 alone. Surprisingly, there were almost 25 fold more mast cells present in co-injected mice (Figure 3C). The findings of increased myeloma cells and increased mast cells in co-injected mice were amplified by the experimental setup in which all three groups received the same total number of cells per mouse (2×10^6^), so co-injected mice only received 10^6^ of MM80 and 10^6^ mast cells compared to twice that in each single group. Lastly, given existing observations that mast cells contribute to bone modeling(14), we hypothesized that the mast cell-myeloma cell interaction may contribute to bone destruction. We assessed for bone destruction in the injected femur of mice in each group by CT scan before euthanization. Both groups of mast cell or MM80 alone injected mice showed similar femur cortical appearance, but co-injected mice showed increased distal femoral bone destruction (Figure 3D).

**Fig 3:**
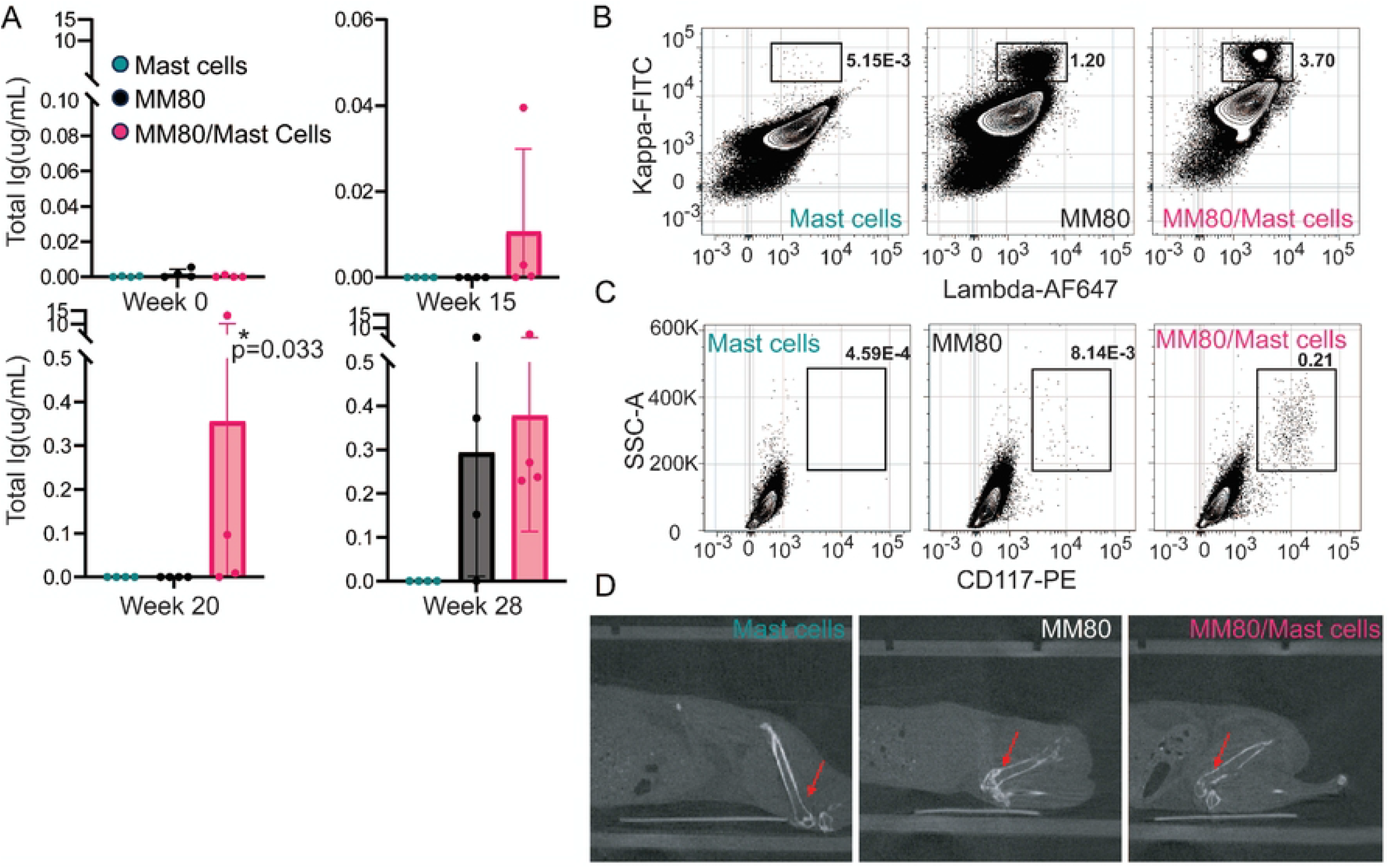
Co-injection of human mast cells with primary CD138 MM cells leads to faster engraftment and bone destruction in the NSG-hIL6 MM PDX model.

NSG-hIL6 were busulfan conditioned and intraosseously injected with human mast cells (green, n=4), CD138+ myeloma cells from the marrow of a newly diagnosed, untreated IgG kappa myeloma patient (MM80) who presented with a pathologic fracture (black, n=4) or CD138+ myeloma cells plus human mast cells (red, n=4). All mice received 2×106 total cells. (A) Serum was collected and analyzed for total human immunoglobulin by ELISA before injection (week 0) and then at 15, 20 and 28 weeks post injection. (B) Representative flow cytometry from bone marrow of each group, gated on singlets, live cells and intracellular kappa vs lambda light chain staining. (C) Representative flow cytometric analysis of bone marrow from each group, gated on singlets, live cells, surface human specific CD117 staining, representative of human mast cells. (D) Representative CT scans of each group of mice at 40 weeks post engraftment depicting the injected femur. Only two mice were alive in the MM80 alone and MM80+mast cell groups at the time of flow cytometry and CT scan analysis, thus statistics were not calculated. p-values determined with one-way ANOVA.

### Mast cells increase myeloma cell survival in vitro through release of secreted factors

After determining that mast cells could decrease engraftment time and increase the percentage of myeloma cells present in the marrow of PDX mice, we sought to determine if human mast cells could increase the longevity of myeloma in cell culture conditions. We plated 2×10^6^ mononuclear cells from three different patient samples, 1 multiply relapsed (MM90) and 2 new diagnoses (MM93, MM80), individually or with mast cells (co-culture was controlled for cell number, 10^6^ of each cell type) for 5 days. Mast cells alone were used as a control. Percentage of live myeloma cells was determined by flow cytometry for intracellular light chain positive cells. All three samples showed significantly increased survival in co-culture conditions (Figure 4A,B). We also sought to determine if the observed increased myeloma survival was due to a contact phenomenon between mast cells and mononuclear cells from the patient or was due to secreted mediators. Using MM80, we put 10^6^ mononuclear cells into a tissue culture plate, placed a tissue insert cup on top and then added 10^6^ mast cells into the cup. This experimental setup prevents contact between cells but allows for secreted factors to flow freely between the compartments. Co-culture of MM80 with mast cells showed the same increased survival as without the tissue culture insert indicating that secreted factors were responsible for the observed improvement in myeloma viability (Figure 4B).

**Fig 4:**
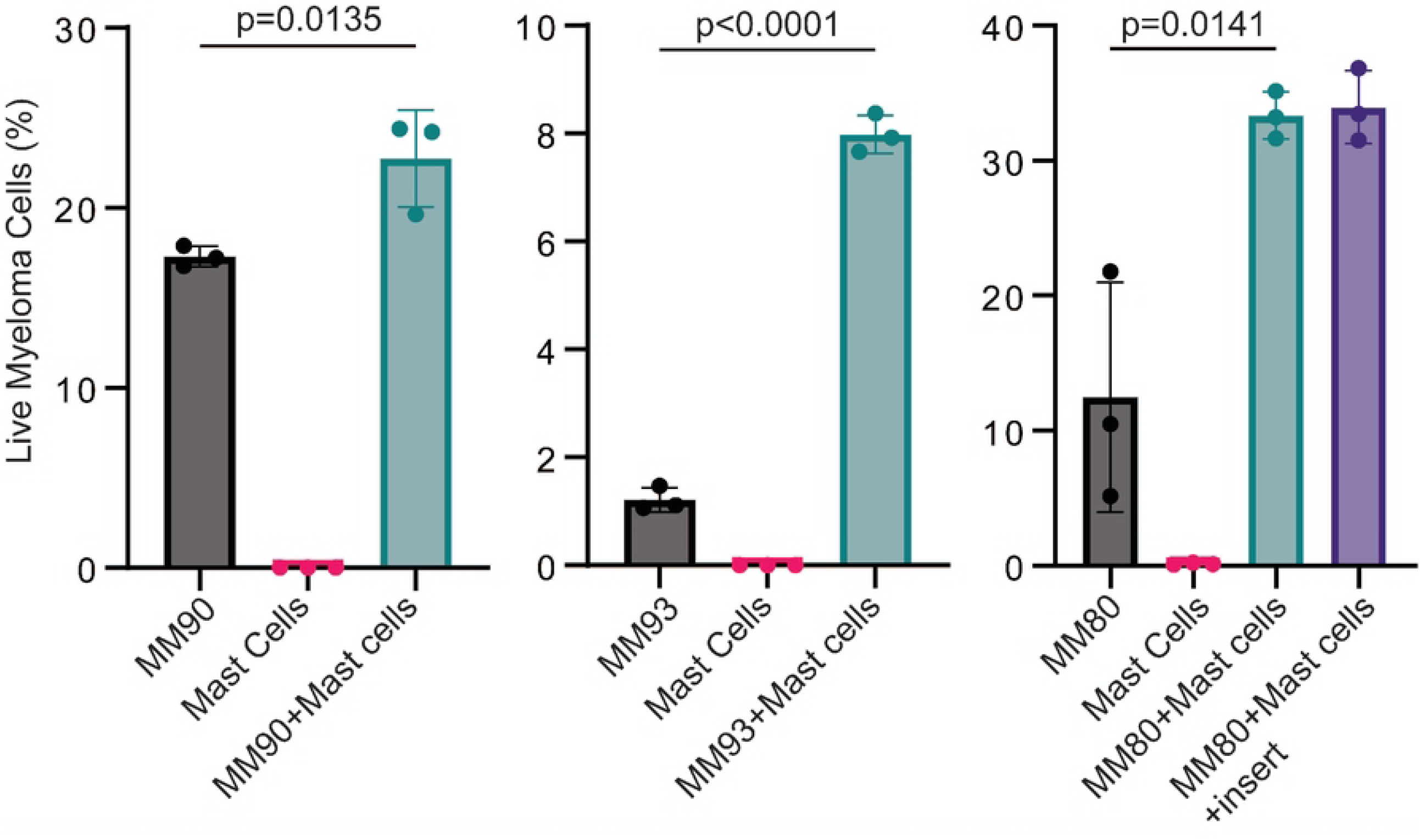
Mast Cells increase survival of primary MM cells ex vivo through secreted factors.

(A,B) Mononuclear cells from three frozen multiple myeloma patient bone marrow aspirates were plated in vitro alone (black) or in the presence of human mast cells. The percentage of live myeloma cells compared to the total live cell fraction was assessed after 72 hours of co-culture by flow cytometry. (B, purple) Additionally, with one myeloma sample (MM80), mononuclear cells were plated in a dish, a tissue culture insert placed into the well and human mast cells placed into the tissue culture insert (cup). Again, the percentage of live myeloma cells was assessed by flow cytometry. Human mast cells cultured alone were used as negative control (pink). p-values determined by one-way ANOVA.

### Ten candidate cytokines were identified from mast cell supernatants

The next step was to identify the mast cell secreted factors responsible for increased myeloma survival. Using acellular supernatants from the MM90 co-culture experiment (Figure 4A), we performed a proteome profiler array looking at 109 potential human cytokines. Of the 109, 40 cytokines showed detectable levels of secretion (Figure 5A). From the 40 detected cytokines, 10 showed expression in mast cell containing supernatants but not in MM90 mononuclear fractions alone (Figure 5B). These cytokines were DPPIV, uPAR, Osteopontin, MIP-3b, GDF-15, sCD14, Dkk-1, CXCL5, IL6 and IGFBP-2. IL6 is known to be secreted from mast cells(15) and served as a functional mast cell control in these experiments.

**Fig 5:**
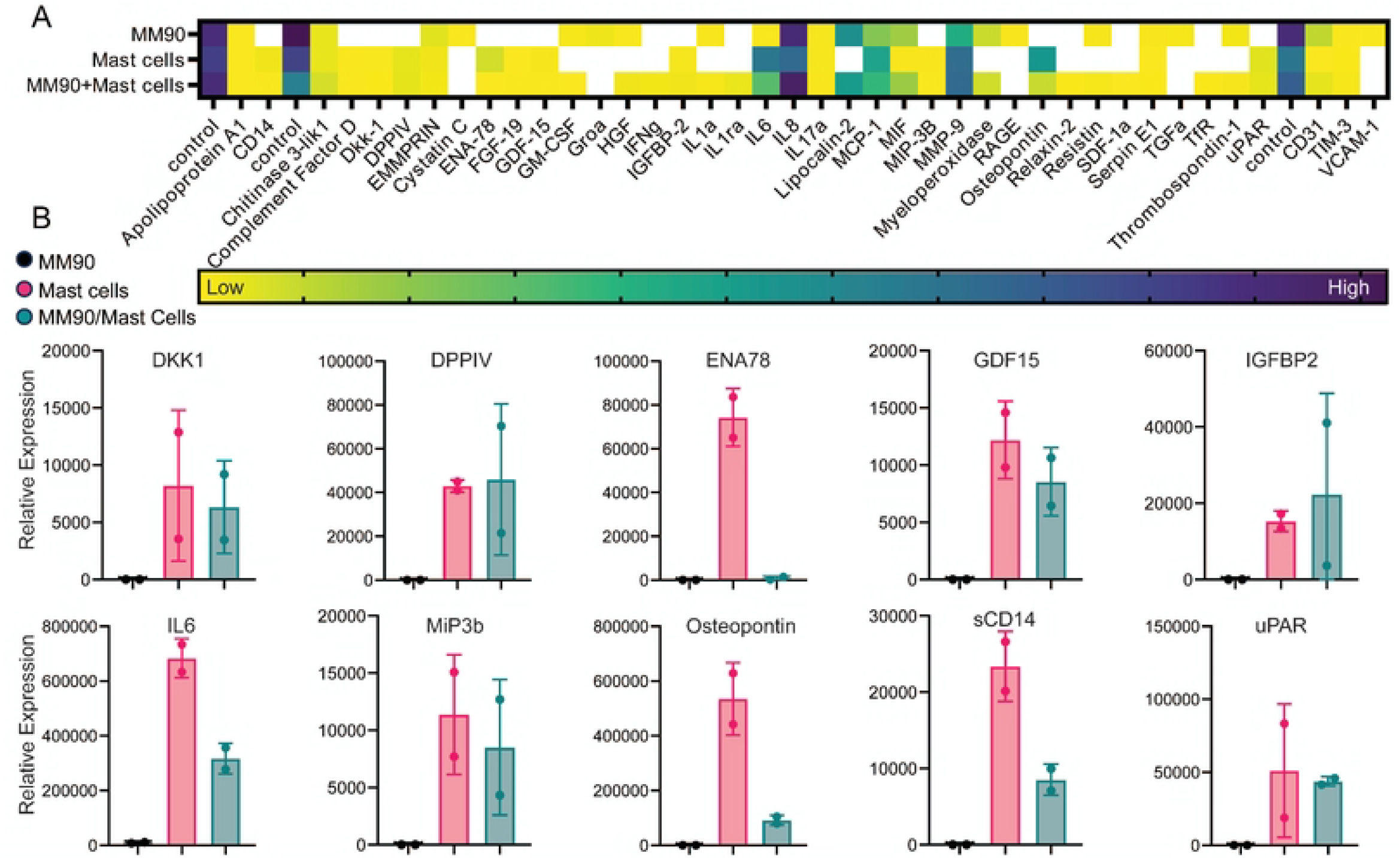
Ten cytokines were increased in mast cell containing supernatants.

Supernatants from cell culture of MM90 with and without mast cells (Figure 4A) were assessed by proteome profiler array for 109 cytokines. (A) 40 cytokines were detected by the array. (B) Cytokines present in mast cell containing conditions but not in mononuclear bone marrow cells alone were graphed. Total plated cell count was maintained at 1e6 cells per condition (MM90 cells co-cultured with mast cells contained 500,000 of each cell type) which is reflected in the overall lower cytokine expression during co-culture compared to mast cells alone. No statistical tests were performed due to only 2 replicates per condition per cytokine, a limitation of the proteome profiler array.

### *Mast cell secreted cytokines support myeloma cell survival* in vitro

We then took MM80 mononuclear cells and cultured them with each of the 10 cytokines identified by the profiler array or all the cytokines together. We chose a high dose of cytokines, 200ng/mL, to maximize the potential effect. A more expansive dose titration of all combinations of cytokines was not feasible. Of the 10 cytokines, 5 singly showed no effect (supplemental figure 4), but sCD14, Dkk-1, IGFBP-2, MIP-3b and uPAR singly showed a significant increase in myeloma cell survival after 5 days of *in vitro* cell culture. However, when all 10 cytokines were added together the largest survival increase was noted, doubling the amount of surviving myeloma cells (Figure 6). We conclude that mast cell secreted cytokines can prolong myeloma cell survival, and multiple cytokines contribute to this process.

**Fig 6:**
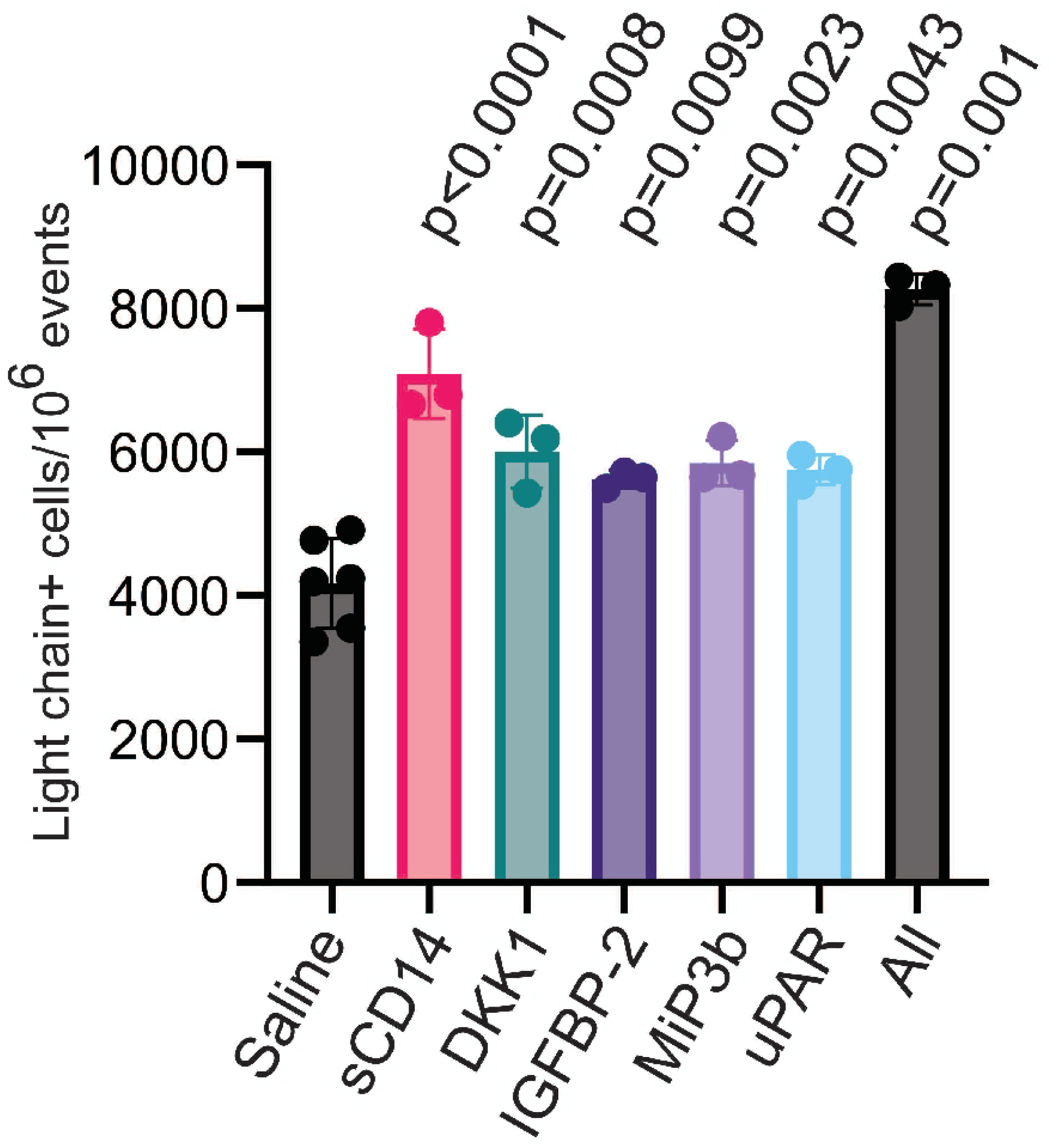
Mast cell secreted cytokines support myeloma cell survival in vitro.

Each of the 10 cytokines from the proteome profiler array (Figure 5B) or all together were added to thawed MM80 mononuclear cells in vitro at a dose of 200ng/mL and cultured for 5 days. Live, singlet, CD3-, CD20-, intracellular kappa light chain+ cells were counted by flow cytometry. Of the 10 cytokines, 5 showed a survival advantage as a single cytokine but all 10 together showed the largest quantitative effect, essentially doubling the surviving cells after 5 days. p-values determined using one-way ANOVA.

## Discussion

This study set out to expand upon an observation that human mast cells were overrepresented in the bone marrow of human myeloma engrafted PDX NSG-hIL6 mice. We first appreciated this phenomenon in the marrow of a single myeloma engrafted mouse on scRNA sequencing. To determine whether present more generally, we assessed for and found mast cells in 5 different xenografts from 5 different donors, suggesting a more widespread phenomenon. We did not select myeloma samples based on genetic or risk-based approaches but rather on the availability of high burden myeloma tissue which engrafts more reliably. Myeloma is a wide and varied disease with differing clinical and molecular presentations, and thus a much larger sample size is needed to determine true generalizability of mast cell support of myeloma survival and contribution to bone lesions. Because of the length and cost of PDX experiments, it is not feasible to use this approach to look at large cohorts of patient samples to determine whether this mast cell phenomenon is more prevalent in some presentations over others. Advanced methodologies such as spatial transcriptomics and bone marrow extracellular fluid proteomics may be helpful to assess large cohorts of patient samples to determine relative involvement of mast cells and level of secreted cytokines but are beyond the scope of this proof of concept study.

Prior studies have expounded upon the correlation of mast cell density in the bone marrow of myeloma patients with poor prognoses.(5) However, we note a discrepancy between our observations in the PDX system regarding the presence of mast cells and their absence in older publications of scRNAseq on human myeloma marrow.(16) We point out that earlier studies only looked at small numbers of cells, 10-20,000, in which the small number of mast cells could be easily lost. Secondly, mast cells are sticky and full of bioactive granules that contribute to low quality RNA and therefore dropout. Despite these limitations, newer studies looking at more cells with deeper sequencing are indeed uncovering mast cell populations in myeloma patient marrow.(17) Though small, they align with findings from prior histological studies that tie mast cell density to ISS staging and prognosis.

In these histology-based studies, mast cells are counted on the order of 1-10 per 0.0625 mm^2^,(5) which is less than 0.1% of the cellularity of the bone marrow. Even with these low numbers of cells, correlations between mast cells, angiogenesis and bone destruction have been proposed.(5) Our *in vitro* data suggests that mast cell-mediated myeloma survival is not a result of angiogenesis alone given that our culture is 2D and absent vascularization. Moreover, classic angiogenic factors, such as VEGF were assayed but did not show expression in our proteome array. We did not perform CD138 isolation of myeloma patient samples before mast cell co-culture due to poor *in vitro* survival after thawing and subsequent column purification. Thus, it is not clear whether mast cells or their cytokines act directly on myeloma cells or act on additional microenvironmental cells that then affect myeloma survival. Future studies will be needed to address this gap.

Regarding bone destruction, we CT scanned the xenografted femur of NSG-hIL6 mice with mast cells only, CD138 selected myeloma cells only or both (Figure 3D) and noted bone destruction in co-injected femurs and not in the other groups. Our proteome array did not include RANKL which is one of the major factors involved in bone destruction in myeloma.(18) However, RANKL cannot be the only factor involved in bone destruction due to myeloma, because myeloma patients with aggressive bone disease on potent anti-RANKL therapies, such as denosumab, continue to develop osteolytic lesions. Not to mention, other non-RANKL targeted bone strengthening therapies, such as zoledronic acid, are equally effective and do not affect RANKL levels.(19)

Of the 5 factors we identified that support myeloma survival, Dkk-1(20) and IGFBP-2(21) have been associated with bone destruction. Dkk-1(22), IGFBP-2(23) and uPAR(24) have also been associated with myeloma progression. IGFBP-2 is required for osteoblast differentiation(25), but high serum levels are found to associate with decreased bone density.(26) We hypothesize that mast cell secreted factors work together to support myeloma survival given that the greatest survival effect was noted when all cytokines were added together versus any one alone (Figure 6). Interestingly, IL6 alone did not confer increased survival to myeloma cells *in vitro*. IL6 is thought of as a requisite survival factor for myeloma, yet, in our experience, it is not routine that IL6 can support *in vitro* growth or survival of myeloma by itself. One explanation is that IL6 actually supports myeloma survival during division, rather than causing division itself.(27) We did not assess cell division directly, because myeloma cell division is difficult to achieve *in vitro*. Moreover, a phase II clinical trial of siltuximab, an anti-IL6 antibody, showed poor results.(28) Taken together all these data suggest that IL6 can be important to myeloma fitness, but it is likely one part of many interactions that confer division and survival advantages.

Methodologically speaking, because primary myeloma cells have low division rates(29) and rapidly die when removed from bone marrow and placed *in vitro*, techniques to increase myeloma *ex vivo* survival would be beneficial. Our data suggest that either mast cells or mast cell cytokines, in particular the 10 identified in this study, lengthen survival time in culturing myeloma *in vitro*. Additionally, this study shows the NSG-hIL6 model as a useful platform to study microenvironmental interactions between myeloma and non-malignant cells through cell mixing experiments. With mast cells, we not only observed an increased survival of myeloma cells, but mast cells were increased when myeloma cells were present *in vivo* (Figure 3C), highlighting a possible symbiotic interplay between the two cell types.

There are several limitations of the current study. Firstly, the mast cells used in experiments were used at concentrations much higher than those observed in the marrow of myeloma patients (∼20% vs <1%) and thus effects seen in our xenograft and culture systems may be of greater magnitude than in patient disease. However, based on the observation that Vκ*-myc cells engrafted faster and survived better with the presence of native mast cells in mice (Figure 2), the biological effects seen in our study may still hold true. We would ideally measure each of the cytokines from the sera of mice and cell culture, but this was not technically feasible in our study. A second major limitation of our study is the sample sizes used. Due to the cost, cell requirements and length of PDX experiments, it is infeasible to expand the PDX system to large cohorts of molecular, genetic or risk-stratified groups. We propose the use of the Myeloma NSG-hIL6 PDX system, as in this study, to determine proof of concept for microenvironmental survival factors and cellular interactions. Subsequently, advanced high resolution spatial transcriptomics and bone marrow proteomics on patient bone marrow core samples may be able to validate findings from PDX experiments in specific larger cohorts, elucidating whether targeting of cell types or microenvironmental proteins is specific to certain myeloma presentations or more generalizable. Thirdly, whether targeting mast cells to improve either myeloma cell killing or prevent bone disease is a viable approach remains unclear. There are not many efficacious mast cell targeted therapies, but inhibiting c-kit with imatinib does appear to reduce bone marrow mast cells and has been used to treat systemic mastocytosis.(30) Because myeloma cells sometimes express c-kit, imatinib was trialed in a single agent phase II study of relapsed/refractory myeloma patients with c-kit positive myeloma cells but did not show any responses.(31) That study, however, had some major limitations. It did not look at the rates of myeloma clinical complications (including lytic bone disease), used imatinib as a single agent and did not consider the mast cell as a target so did not assay the mast cell pool. There may still be potential for mast cell targeted therapies to improve bone strength and possibly even outcomes in myeloma, but larger pre-clinical studies are needed.

## Acknowledgements

The authors would like to thank the UPenn Stem Cell and Xenograft Core and Derek Dopkin for their help with setting up PDX experiments, the Small Animal Imaging Facility for facilitating CT scans, Leif Bergsagel and Marti Chesi for providing Vκ*-myc cells and the UPenn clinical stem cell lab for providing CD34+ stem cells. This study was made possible by funding from the UPenn CTSA KL2 (Hasanali) and Mark Foundation (Allman).

